# First record of the endophytic bacteria of *Deschampsia antarctica* E. Desv. from two distant localities of the maritime Antarctica

**DOI:** 10.1101/586099

**Authors:** O. Podolich, I. Parnikoza, T. Voznyuk, G. Zubova, I. Zaets, N. Miryuta, G. Myryuta, O. Poronnik, I. Kozeretska, V. Kunakh, A.M. Pirttila, N. Kozyrovska

## Abstract

The vascular plant *Deschampsia antarctica* samples were collected for endophytic bacteria study from two regions in the maritime Antarctic 400 km distant from one another: Point Thomas oasis (King George Island) and Argentine Islands (Galindez Island). The endophytes were isolated from roots and leaves of *D. antarctica*, cultivated and identified by using a partial sequencing of the 16S rRNA gene served as a phylomarker. Endophyte isolates from two sites of Galindez Island were represented mainly by *Pseudomonas* species and by *Gammaproteobacteria, Firmicutes* and *Actinobacteria*. The vast majority of the isolates had specific for endophytes cellulase and pectinase activities, however, *Bacillus* spp. did not express both activities. A group-specific PCR screening at the four sites of Galindez Island and two sites of King George Island, indicated *Alphaproteobacteria, Betaproteobacteria, Gammaproteobacteria, Firmicutes, Cytophaga-Flavobacteria* and *Actinobacteria.* Notably, the number of endophytic bacteria taxa was significantly larger in leaves than in roots of plants.

## Introduction

Endophytic microorganisms live inside the asymptomatic host plants as commensals (van Overbeek and Saikkonen 2016) and are known to have beneficial effect on plants such as promoting the plant growth and the plant protection from different biotic and abiotic stresses. Endophytic bacteria are ubiquitous inside of all plant species and create together with the host plant a superorganism – a multispecies community that functions as an organizational unit (Podolich et al. 2015). The major source of the endophytes in plants is rhizo- and phyllosphere (Edwards et al. 2015).

The maritime Antarctica presents the global southern limit for a spread of the vascular plants, with only two vascular plants species found, and one of these species is Antarctic hairgrass *Deschampsia antarctica* E. Desv. (*Poaceae*) (Parnikoza et al. 2009; Parnikoza et al. 2017). The adaptation of the Antarctic hairgrass to cold environment could be supported by individual and unique cross-species interactions, *e.g., D. antarctica* has no specific adaptive mechanisms, therefore its survival and colonization of the Antarctica may depend on the interactions with another organisms, like birds (Parnikoza et al. 2018) or bacteria. Moreover, plants, living in extreme environments, can be a rich source of associated bacterial species, beneficial for biotechnological purposes, *e.g.*, as producers of enzymes active under low temperatures (Kuddus 2018). Antarctic plant-associated bacteria have been recovered from mosses (Park et al. 2013), as well as endophytic fungi were found in *Deschampsia* (Santiago et al. 2016). However, there is no information about the prokaryotic endophytes of *D. antarctica* from the different maritime Antarctica.

## Material and Methods

Sampling was conducted in two regions of the maritime Antarctica, separated by a distance of 400 km from one another: Point Thomas oasis, King George Island, South Shetland Islands, and the Argentine Islands during the austral summer season 2014, and 2017/18. We collected green plants with a near root substrata (leptosols) – three specimens of *D. antarctica* plants from each locality (Supplement Table 1), packed them in sterile plastic boxes and transported to the laboratory. Endophytic bacteria were isolated from two locations of Galindez Island, namely, Karpaty Ridge and Metheo Point. For isolation of endophytic bacteria, *D. antarctica* plants were surface-sterilized in 70% ethanol for 1 min and in 6% calcium hypochlorite for 20 min and rinsed three times for 5 min in sterile distilled water. The last washing was controlled on dissemination by inoculation of DW on nutrient agar and no microbial growth was observed. The plant material was crushed in mortar with a pestle, serially diluted and cultivated on KB, LB and M9 agar media. Bacterial DNA isolation was performed with innuSPEED bacteria/fungi DNA isolation kit (Analytik Jena AG). Endophytic bacterial isolates were identified by a PCR amplification, using standard primers 27F and 1492R (Fredriksson et al. 2013). The PCR products were sequenced by the Sanger method using Big Dye Terminator Sequencing Standard Kit v3.1 (Applied Biosystems, USA) and apparatus 3130 Genetic Analyser (Applied Biosystems). The 16S rDNA sequences were binned by BLASTN programs search through the NCBI (USA). GenBank accession numbers: MG916945-MG916956. Neighbour-joining phylogenetic tree were conducted in MEGA 7. (Supplement, Fig.1).

The endophytic isolates were examined primarily for cellulolytic (Wood 1981) and pectinolytic (Starr et al. 1977) activities by plate assays.

For group-specific bacterial amplification, total DNA was isolated from the surface-sterilized roots and leaves of *D*. *antarctica* plants from all localities, using Power Plant DNA isolation kit (MoBio). The isolated DNA (100 ng) was amplified with group-specific bacterial primers: Alf28f/Alf684r for *Alphaproteobacteria*, Beta359f/Beta682r for *Betaproteobacteria*, Gamma395f/Gamma871r for *Gammaproteobacteria*, Firm350f/Firm814r for *Firmicutes* (Mühling et al. 2008), ACT235f/ACT878r for *Actinobacteria*, CF315f/ CF967r (Stach et al. 2003), *Cytophaga – Flavobacterium* (Xihan et al. 2008), and the PCR products were separated in a 2.0% agarose gel.

## Results and Discussion

Identification and phylogenetic analysis (Supplement, Fig.1) of the endophytic bacterial isolates from the roots of *D. antarctica* showed that representatives of the *Pseudomonas* genus were the most common in the plant interiors of both localities (Supplement, Table 2). Interestingly, all gram-negative endophytic isolates were represented exclusively by *Gammaproteobacteria*. The vast majority of them had cellulase or pectinase activities; *P. migulae, P. rhodesiae, P. orientalis* and *P. antarctica* had both activities, but *P. graminis* and *P. fluorescens* did not exhibit such traits. It is known that bacterial genus *Pseudomonas* has appeared frequently in different Antarctic environments (Vásquez-Ponce et al. 2018). Also, the genus *Pseudomonas* is tightly associated as endophytes with circumpolar grass *Deschampsia flexuosa* (L.) Trin., growing in subarctic Aeolian sand dune area (Poosakkannu et al. 2014). Thus, the results indicate that bacteria of the genus *Pseudomonas* frequently inhabit extremely cold environment.

Gram-positive bacteria were present only in plant samples from Metheo Point, and they belonged to the phyla *Firmicutes* and *Actinobacteria*. Remarkably, isolated *B. subtilis* and *B. pumilis* did not express both activities, although it is known that cellulose activity is common for *Bacillus* spp., occupying different mainland econiches (Gupta et al. 2015), including endophytic *Bacillus* and *Paenibacillus* strains exhibited both cellulase and pectinase activities (Zhao et al. 2015).

Results of a group-specific PCR assay, which was used for characterization of both culturable and unculturable endophytes of the hairgrass, represented in Supplement, Fig. 2. For all studied regions, cultivation-independent approach showed more taxons in endophyte community structures than a culture-based one: *Proteobacteria (Alpha-, Beta-, Gammaproteobacteria), Firmicutes, Cytophaga-Flavobacteria* and *Actinobacteria*. There was some difference in the studied groups of endophytic bacteria between the roots and leaves. The number of taxa of studied endophytic bacteria was wider in leaves than in roots of the plants from the two sites of Galindez Island (Karpaty Ridge and Magnit Cape) and one site of King George Island (Point Thomas, Puchalski grave). Such difference was especially noticeable in plants from the site of Magnit Cape (Galindez Island). It could be explained by specific conditions in substrates (Zaets et al. 2012), *e.g*., the high content of ions of trace elements in soils in some localities of the maritime Antarctic. Homogeneity of endophytic community structure was observed for the developed *D. antarctica* cenoses within distant locations, *e.g*., the plants from King George Island, Point Thomas, Puchalski grave and Galindez Island, Cemetry Ridge had similar endophytic communities in both roots and leaves.

## Conclusion

Summarizing, we may conclude that in isolates from *D. antarctica* most abundant genus was *Pseudomonas.* Some of isolated cultures had pectinase and cellulase activities. Remarkably, the representatives of *Firmicutes* did not possess cellulase and pectinase activities, as compared to homologous species in another parts of the world. This may mean that horizontal gene transfer events could take place rarer in the maritime Antarctic than on the mainland. In some regions, the culture independent approach indicated a higher number of bacterial taxons in leaves than in roots, which may be explained by extreme Antarctic environmental conditions.

## Acknowledgments

The fieldwork was supported by the National Antarctic Scientific Centre of the Ministry of Science of Ukraine during the 18th Ukrainian Antarctic expeditions and approved by Department of Antarctic study of Institute of Biochemistry and Biophysics PAS. We would like to thank Dr Papitashvili for their help in the expedition preparation and A. Berezkina for her valuable help in the map preparation, and Mag. M. Wierzgon for the samples collection.

## Compliance with Ethical Standards

This study was not accompanied by the emergence of potential conflicts of interest and did not include Human Participants or Animals.

---

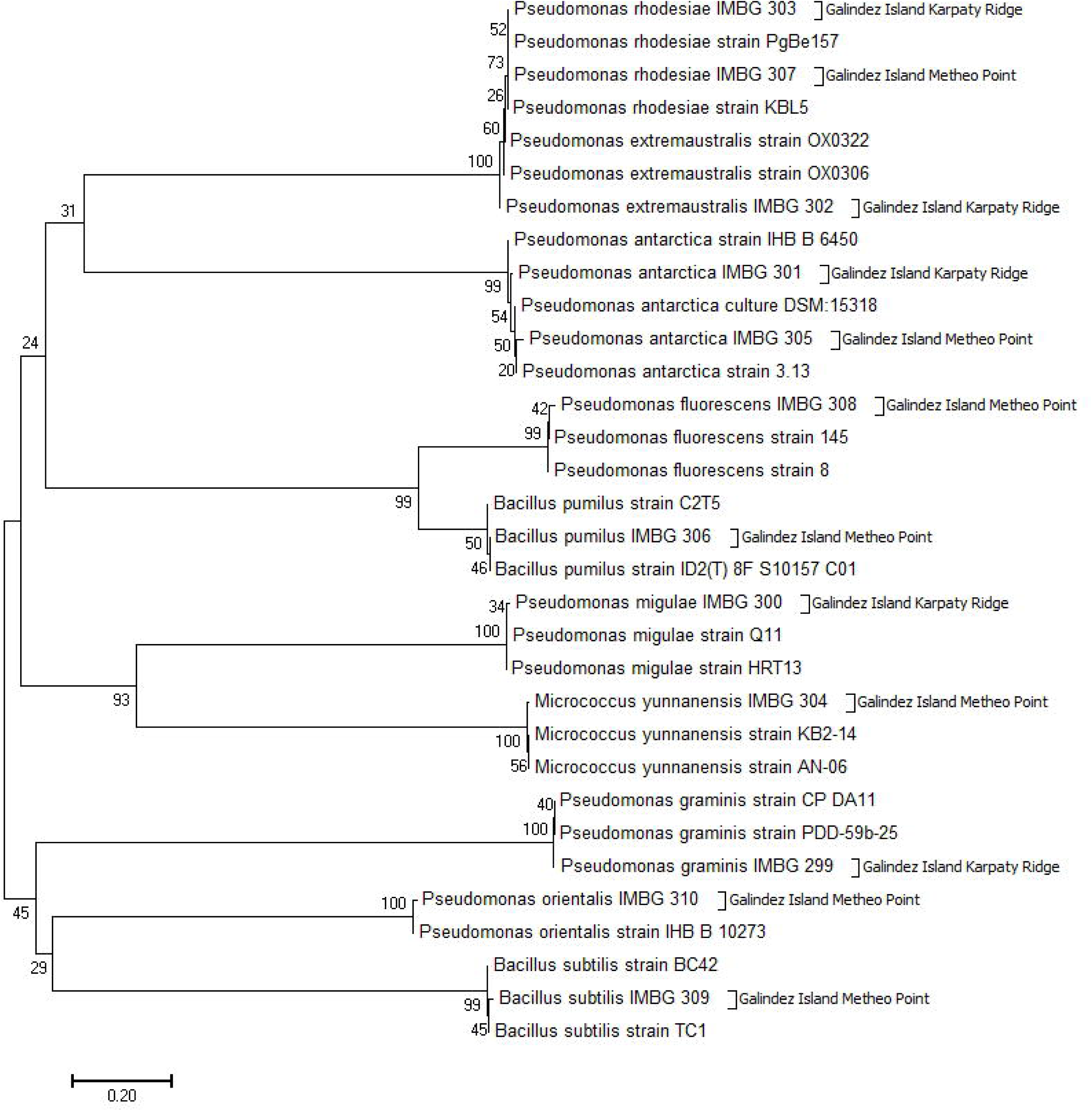

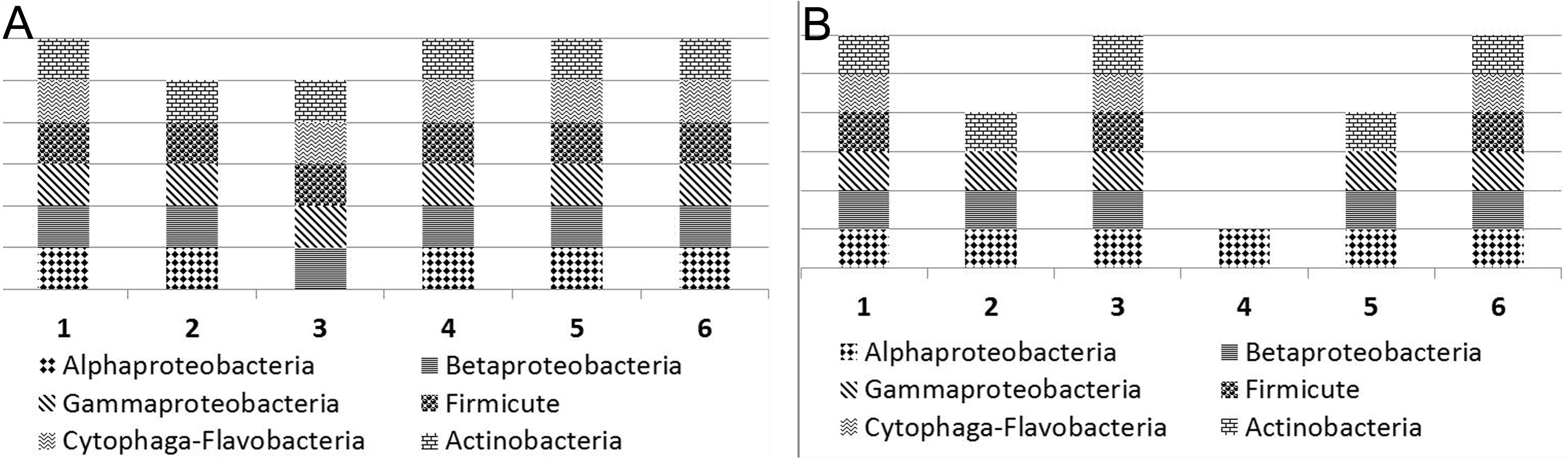

## References

Fredriksson NJ, Hermansson M, Wilén BM (2013) The choice of PCR primers has great impact on assessments of bacterial community diversity and dynamics in a wastewater treatment plant. PLoS One 8(10):e76431

Gupta M, Sharma M, Singh S, Gupta P, Bajaj BK (2015) Enahnced production of cellulase from *Bacillus Licheniformis* K-3 with potential saccharification of rice straw. Energy Technol 3:216–224

Kuddus M (2018) Cold-active enzymes in food biotechnology: An updated mini review. Journal of Appl Biol Biotechnol 6(3):58–63

Mühling M, Woolven-Allen J, Murrell JC, Joint I (2008) Improved group-specific PCR primers for denaturing gradient gel electrophoresis analysis of the genetic diversity of complex microbial communities. The ISME Journal 2:379–392

Park M, Lee H, Hong SG, Kim OS (2013) Endophytic bacterial diversity of an Antarctic moss, *Sanionia uncinata*. Antarctic Science 25(1):51–54

Parnikoza I, Convey P, Dykyy I, Trokhymets V, Milinevsky G, Inozemtseva D, Kozeretska I (2009) Current status of the Antarctic herb tundra formation in the central Argentine Islands. Global Change Biol 15:1685–1693

Parnikoza I., Abakumov E., Korsun S., Klymenko I., Netsyk M., Kudinova A., Kozeretska I. (2017) Soils of the Argentine Islands, Antarctica: Diversity and Characteristics. Polarforschung 86 (2):83–96, doi:10.2312/polarforschung.86.2.83

Parnikoza I, Rozhok A, Convey P, Veselski M, Esefeld J, Ochyra R, Mustafa O, Braun C, Peter HU, Smykla J, Kunakh V, Kozeretska I (2018) Spread of Antarctic vegetation by the kelp gull: comparison of two maritime Antarctic regions. Polar Biology. doi.org/10.1007/s00300-018-2274-9

Podolich O, Ardanov P, Zaets I, Pirttilä AM, Kozyrovska N (2015) Reviving of the endophytic bacterial community as a putative mechanism of plant resistance. Plant Soil 388:367–377

Poosakkannu A, Nissinen R, Kytöviita MM (2014) Culturable endophytic microbial communities in the circumpolar grass, *Deschampsia flexuosa* in a sub-Arctic inland primary succession are habitat and growth stage specific.Environmental Microbiology Reports doi:10.1111/1758-2229.12195

Santiago IF, Rosa CA, Rosa LH (2016) Endophytic symbiont yeasts associated with the Antarctic angiosperms *Deschampsia antarctica* and *Colobanthus quitensis*. Polar Biol doi:10.1007/s00300-016-1940-z

Stach JEM, Maldonado LA, Ward AC, Goodfellow M, Bull AT (2003) New primers for the class *Actinobacteria*: application to marine and terrestrial environments. Environ Microbiol 5:828–841

Starr MP, Chatterjee AK, Starr PB, Buhanan GE (1977) Enzymatic degradation of polygalacturonic acid by *Yersinia* and *Klebsiella* species in relation to clinical laboratory procedures. J Clin Microbiol 6:379–386

van Overbeek LS, Saikkonen K (2016) Impact of bacterial–fungal interactions on the colonization of the endosphere. Trends Plant Sci 21:230–242

Wood PJ (1981) The use of dye-polysaccharide interactions in β-D-glucanase assay. Carbohydr Res 94:19–23

Xihan Ch, Zeng Y, Jiao N (2008) Characterization of *Cytophaga-Flavobacteria* community structure in the bering sea by cluster-specific 16S rRNA gene amplification analysis. J Microbiol Biotechnol 18(2):194–198

Zaets I., Kozyrovska N. (2012) Heavy Metal Resistance in Plants: A Putative Role of Endophytic Bacteria. Toxicity of Heavy Metals to Legumes and Bioremediation. Editor Zaidi et al. Springer Verlag Wien 203-217

Zhao L, Xu Y, Lai XH, Shan C, Deng Z, Ji Y (2015) Screening and characterization of endophytic *Bacillus* and *Paenibacillus* strains from medicinal plant *Lonicera japonica* for use as potential plant growth promoters. Braz J Microbiol 46(4):977–89.

