## Supplemental tables for "First record of the endophytic bacteria of *Deschampsia antarctica* E. Desv. from two distant localities of the maritime Antarctica"

**Supplement tables**

Table 1. Characteristics of sampled points of *Deschampsia antarctica* E. Desv. on Point Thomas, King George Island and Galindez Island, Argentine Islands

| # | Sample location | Coordinates | *Short description* |
| --- | --- | --- | --- |
| 1. | King George Island, Point Thomas, Puchalski grave | 62° 9.811' S, 58° 28.146' W | On the top of Puchalski grave, Total Vegatation Cover (TVC) 90%, *D. antarc­tica* 50%, *Colobantus quitensis* – 1 %, Bryophytes – 10 %, *Usnea antarctica* – 5-10%, *Ochrolechia* sp. –5%, 28 m.a.s.l. |
| 2. | King George Island, Point Thomas | 62° 9.989' S, 58° 28.097' W | Area with gravel, initial stage of vegetation colonization, TVC 1%, small cover of *D. antarc­tica* 0,5%, *C. quitensis* – 0,4%, Bryophytes 0,1 %, *D. antarctica*, 50 m a.s.l. |
| 3. | Galindez Island, Metheo Point | 65° 14.687' S, 64° 15.348' W | On rocky coast of Marina Point near Meteorological station, TVC 1 %, *D. antarctica* 0,5 %, *Sanionia* sp*.* 0,5 %, gravel, 13 m.a.s.l. |
| 4. | Galindez Island, Magnit Cape | 65° 14.704' S, 64° 15.155' W | Top of the coastal rock, TVC 5-25 %, *D. antarctica* 4-24 %, Bryophytes 1 %, on limpet shells, 6 m.a.s.l. |
| 5. | Galindez Island, Karpaty Ridge | 65° 14.766' S, 64° 14.951'W | Karpaty Ridge, N slope of central part of the ridge, *Polytrychium strictum* Bridel moss bank, TVC 80 % with incorpo­ration of *Sanionia georgicouncinata* (Müll. Hal.) Ochyra, 65°14.768' S, 64°14.959' W, 17 m.a.s.l. |
| 6. | Galindez Island, Cemetry Ridge | 65° 14.770' S, 64° 14.874' W | Top of the Cemetery Ridge near VLF, TVC 5-40 %, *D. antarc­tica* 4-30 %, bryophytes 1-10 %, limpets, gravel, 17 m.a.s.l. |

Table 2. Endophytic bacteria isolates of *Deshcampsia antarctica* from two localities of Galindez Island

| **Endophytic isolate** | **Accession number** | **Tentative phylogenetic group** | **Localities** | **Enzymatic activity** | |
| --- | --- | --- | --- | --- | --- |
|  |  |  |  | **Pectinase** | **Cellulase** |
| *Pseudomonas graminis* IMBG 299 | MG916945 | *Gammaproteobacteria* | Galindez Island, Karpaty Ridge | - | - |
| *Pseudomonas migulae* IMBG 300 | MG916946 | *Gammaproteobacteria* | Galindez Island, Karpaty Ridge | + | + |
| *Pseudomonas antarctica* IMBG 301 | MG916947 | *Gammaproteobacteria* | Galindez Island, Karpaty Ridge | - | + |
| *Pseudomonas extremaustralis* IMBG 302 | MG916948 | *Gammaproteobacteria* | Galindez Island, Karpaty Ridge | - | + |
| *Pseudomonas rhodesiae* IMBG 303 | MG916949 | *Gammaproteobacteria* | Galindez Island, Karpaty Ridge | + | - |
| *Micrococcus yunnanensis* IMBG 304 | MG916950 | *Actinobacteria* | Galindez Island, Metheo Point | - | - |
| *Pseudomonas antarctica* IMBG 305 | MG916951 | *Gammaproteobacteria* | Galindez Island, Metheo Point | + | + |
| *Bacillus pumilus* IMBG 306 | MG916952 | *Firmicutes* | Galindez Island, Metheo Point | - | - |
| *Pseudomonas rhodesiae* IMBG 307 | MG916953 | *Gammaproteobacteria* | Galindez Island, Metheo Point | + | + |
| *Pseudomonas fluorescens* IMBG 308 | MG916954 | *Gammaproteobacteria* | Galindez Island, Metheo Point | - | - |
| *Bacillus subtilis* IMBG 309 | MG916955 | *Firmicutes* | Galindez Island, Metheo Point | - | - |
| *Pseudomonas orientalis* IMBG 310 | MG916956 | *Gammaproteobacteria* | Galindez Island, Metheo Point | + | + |

**Supplement Figure captures**

Supplement Fig 1. Neighbour-joining phylogenetic tree based on 16S rRNA gene sequences of endophytic bacteria showing the relationship between the endophytes sequences with closest type strain sequences. The percentage of replicate trees in which the associated taxa clustered together in the bootstrap test (500 replicates) are shown next to the branches. The evolutionary distances were computed using the Maximum Composite Likelihood method and are in the units of the number of base substitutions per site. Evolutionary analyses were conducted in MEGA7.

Supplement Fig. 2. Characterization of number of taxa for studied groups of endophytic bacteria in *Deschampsia antarctica* E. Desv. with group-specific PCR. A – leaves DNA samples; B – roots DNA samples. 1. King George Island, Point Thomas, Puchalski grave; 2. King George Island, Point Thomas Near Ecology Glacier; 3. Galindez Island, Metheo Point; 4. Galindez Island, Magnit Cape; 5. Galindez Island, Karpaty Ridge; 6. Galindez Island, Cemetry Ridge
